# Human Endometrial Stromal Cells Are Highly Permissive To Productive Infection by Zika Virus

**DOI:** 10.1101/077305

**Authors:** Isabel Pagani, Silvia Ghezzi, Adele Ulisse, Alicia Rubio, Filippo Turrini, Elisabetta Garavaglia, Massimo Candiani, Concetta Castilletti, Giuseppe Ippolito, Guido Poli, Vania Broccoli, Paola Panina-Bordignon, Elisa Vicenzi

**Author notes:** co-first authors (Isabel Pagani, Silvia Ghezzi). co-last authors (Paola Panina-Bordignon, Elisa Vicenzi). Correspondence (P.P.-B.), (E.V.).

## Abstract

Zika virus (ZIKV) is a recently re-emerged flavivirus transmitted to humans by mosquito bites but also from mother to fetus and by sexual intercourse. We here show for the first time that primary human endometrial stromal cells (HESC) are highly permissive to ZIKV infection and support its *in vitro* replication. ZIKV envelope expression was detected in the endoplasmic reticulum whereas double-stranded viral RNA colocalized with vimentin filaments to the perinuclear region. ZIKV productive infection also occurred in the human T-HESC cell line with the induction of interferon-β (IFN-β) and of IFN-stimulated genes. Notably, *in vitro* decidualization of T-HESC with cyclic AMP and progesterone upregulated the cell surface expression of the ZIKV entry co-receptor AXL and boosted ZIKV replication by *ca.* 100-fold. Thus, endometrial stromal cells, particularly if decidualized, likely represent a crucial cell target of sexual virus transmission and a relevant source of ZIKV spreading to placental trophoblasts during pregnancy.

**AUTHOR SUMMARY:** Infection by Zika virus (ZIKV), a flavivirus transmitted to humans by mosquito bites, has recently emerged as an important cause of neurological lesions in the fetal brain as women who become infected by ZIKV during pregnancy can transmit the virus to their fetus. In addition, routes of ZIKV transmission independent of mosquito bites have been also identified and include sexual transmission from both infected men and women to their partners, an aspect bearing great societal implications for ZIKV spread. These observations highlight the importance of the female reproductive tract in the establishment and/or spreading of the infection. In this regard, the endometrium is a highly dynamic tissue undergoing major histological changes during the menstrual cycle under the coordinated action of sexual hormones. In particular, progesterone drives the differentiation of human endometrial stromal cells towards decidualization, a process that is critical for fetal trophoblast invasion and placenta formation. We here report for the first time that both primary and immortalized human endometrial stromal cells are highly permissive to ZIKV infection and replication, particularly when *in vitro* decidualized by progesterone, suggesting that these cells could significantly contribute to vertical ZIKV transmission in utero during pregnancy but also to horizontal transmission by the sexual route.

## INTRODUCTION

Zika Virus (ZIKV) is a member of the *Flaviviridae* family transmitted to humans by mosquito bites of the *Aedes* species [1]. The virus has been first isolated from the blood of a febrile monkey in 1947 in the Zika forest of Uganda [2]; however, its potential as a human pathogen was underestimated for almost 60 years until 2013 when an unsual outbreak of ZIKV-related Guillain-Barré syndrome emerged in French Polynesia [3]. A global health emergency was triggered at the end of 2015 by the obervation of an increased incidence of microcephaly was associated with the temporal and geographic distribution of ZIKV infection in the North East Brazil [4]. Increasing evidence now clearly supports a cause-effect relationship between congenital ZIKV transmission and increased frequency of mild to severe neuropathologies including microcephaly [5,6]. ZIKV was detected in the amniotic fluid of pregnant women [7] suggesting that the placenta might be permissive to virus passage, a condition that likely occurs during the first trimester of pregnancy [8], although placental cells appear to be protected against ZIKV infection by a constitutive interferon (IFN)-λ1 response [9]. Indeed, two recent studies showed that human primary placental macrophages and trophoblasts were permissive to ZIKV productive infection both *in vitro* [10,11] and in a mice [12]. These studies demonstrates that ZIKV can cross the placental barrier either coming from maternal blood or by an ascending route as recently demonstrated in a mouse model of vaginal ZIKV infection [13].

Vector-independent transmission of ZIKV among humans can occur through the sexual route [14]. Male-to-female transmission has been reported in approximately 15 cases thus far [15]. This hypothesis is also supported by both the detection of higher viral load in semen *vs.* blood and by the persistence of ZIKV in semen for several months after waning of symptoms [16,17,18,19]. More recently, female-to-male sexual transmission of ZIKV infection was also documented [20]. These observations imply a potentially prominent role of the female reproductive tract (FRT) as a site of virus infection and propagation either from and to the male partner during sexual intercourse or to the fetus during pregnancy. All compartments of the FRT, including the endometrium, might contribute to establishing and spreading the initial infection. In addition, it should be taken into consideration the fact that the human endometrium is a highly dynamic tissue undergoing major histological changes during the menstrual cycle under the coordinated action of sexual hormones. Estrogen dominates the proliferative phase of the menstrual cycle, while the post-ovulatory rise of ovarian progesterone drives the differentiation of human endometrial stromal cells (HESC). This process, known as pre-decidualization, is critical for fetal trophoblast invasion and placenta formation and occurs independently of an implanting blastocyst [21].

Therefore, we investigated whether primary HESC or immortalized (T-HESC) cells were permissive to *in vitro* ZIKV infection and replication. Indeed, ZIKV productively infected both HESC and T-HESC, whereas *in vitro* decidualization of the cell line (dT-HESC) increased both the expression of putative ZIKV entry co-receptor AXL and the levels of productive infection *vs.* unstimulated cells. Thus, our results fully support the hypothesis of a relevant role of the endometrium for both sexual ZIKV transmission and *in utero* virus transmission to the fetus during pregnancy.

## RESULTS

### ZIKV infection of primary HESC

Primary HESC were isolated from endometrial biopsies and incubated with either the reference African MR766 or contemporary INMI-1 strains at the multiplicity of infection (MOI) of 10 after reaching cell confluency (days 3-4). Viral growth was firstly analyzed by indirect immunofluorescence (IF) analysis using either an anti-dsRNA monoclonal antibody (mAb) or an anti-panflavivirus envelope (E) protein mAb. Subcellular distributions of both viral RNA and E protein were observed 72 h after infection of HESC (**Figure 1A**). The percentage of cells infected with the MR766 strain, as measured by both anti-dsRNA and anti-flavi E mAb staining, was *ca.* 80% whereas the proportion of cells infected with the INMI-1 strain was 10-fold lower (ca. 7%). Progeny infectious virion production was measured in HESC culture supernatants using a standard plaque-forming assay (PFA) in Vero cells (**Figure 1B**) whereas absolute quantification of viral NS5 RNA was determined by droplet digital (dd) PCR (**Figure 1C**). Both viral strains replicated efficiently in primary HESC established from 8 independent donors although the more recent INMI-1 strain was *ca.* 1 log_10_ less efficient than the MR766 strain, confirming the IF results.

**Figure 1.**
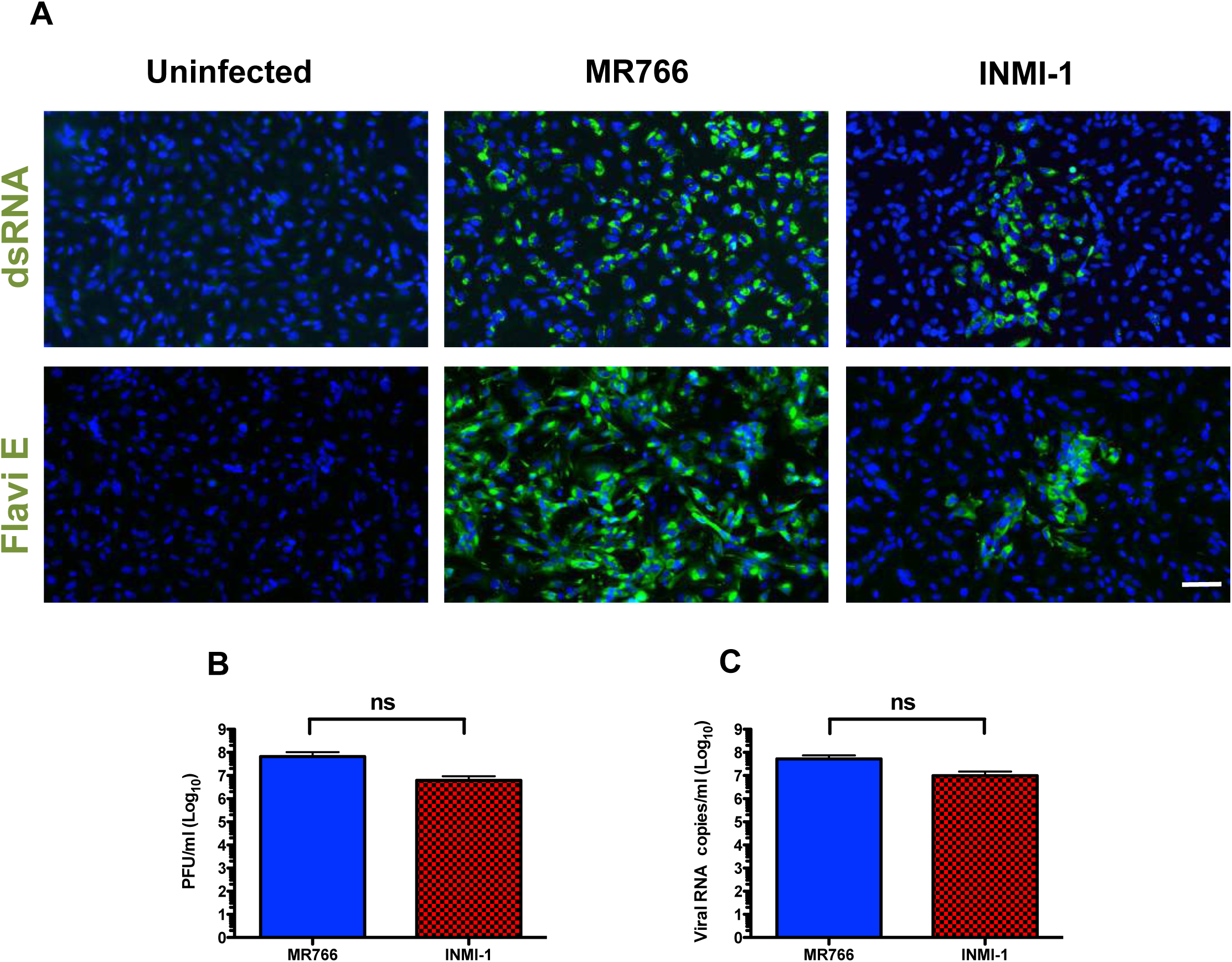
Primary HESC are permissive to ZIKV productive infection. **(A)** Immunostaining for dsRNA and Flavi E of primary HESC infected with MR766 and INMI-1 ZIKV strains 72 h post-infection; Hoechst was used to stain nuclei. Scale bar: 20 µm. **(B)** Viral titers in HESC supernatant harvested 3 days after infection as determined by a PFA in VERO cells. Mean ± SEM of 8 independent donors in duplicate cultures. Ns, not significant. **(C)** Viral RNA quantification in HESC supernatants harvested 3 days post-infection as determined by ddPCR. Mean ± SEM of 8 independent donors.

### Subcellular localization of ZIKV in HESC

The flavivirus life cycle occurs in the cytoplasm mostly in association with intracellular membranes of the endoplasmic reticulum (ER) [22]. Therefore, we analyzed whether ZIKV E protein co-localized with the ER in infected cells by staining infected and uninfected cells with a mAb directed to calreticulin, a major Ca^2+^-binding protein and chaperone expressed in the lumen of the ER [23]. Calreticulin staining in uninfected cells appeared as perinuclear fibrillar fluorescence (**Figure 2A**) whereas in infected cells its staining expanded in the cytoplasm towards the cell perifery. In addition, dense structures colocalized viral E protein expression and calreticulin in the perinuclear region (**Figure 2A**).

**Figure 2.**
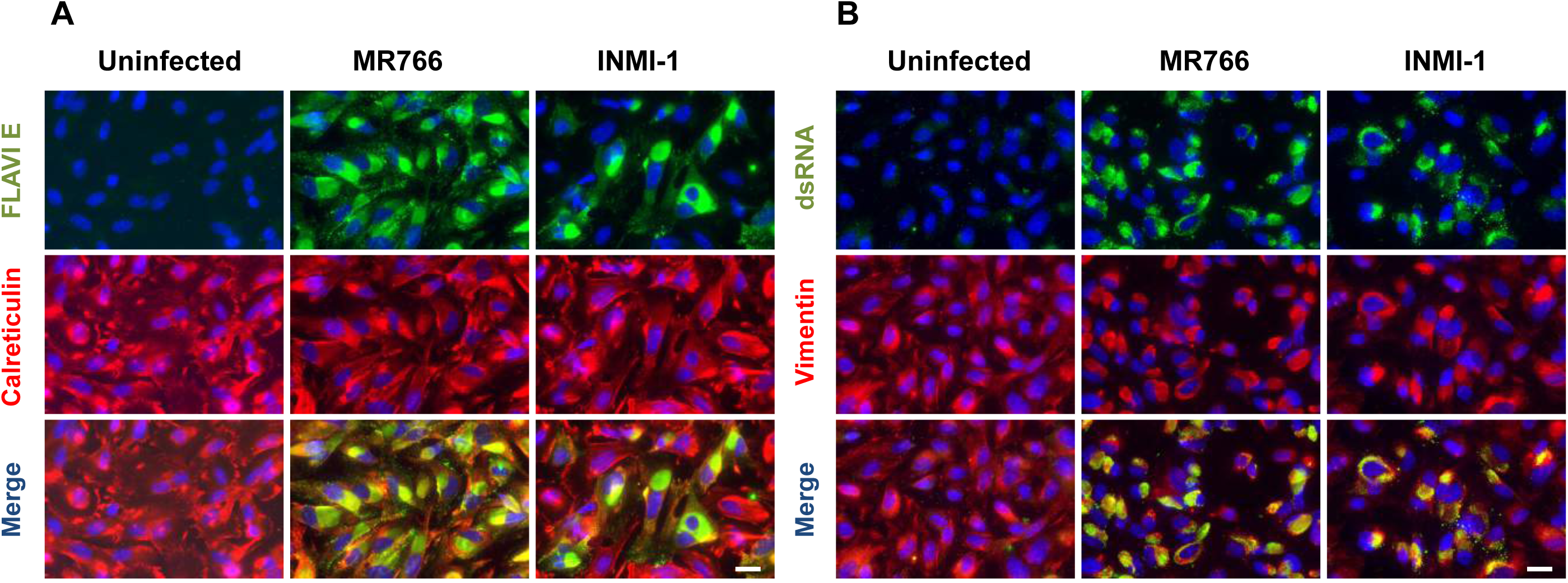
Co-localization of ZIKV with calreticulin and vimentin in HESC. **(A)** Double immunostaining for Flavi E and calreticulin in primary HESC either uninfected or infected with MR766 and INMI-1 at 72 h post-infection; Hoechst was used to stain nuclei. Scale bar: 10 µm. **(B)** Double immunostaining for dsRNA and vimentin in primary HESC either uninfected or infected with MR766 and INMI-1 at 72h post-infection; Hoechst was used to stain nuclei. Scale bar: 10 µm.

We also evaluated whether vimentin, a major component of type III intermediate filaments expressed by HESC [24], colocalized with ZIKV dsRNA. It has been reported previously that vimentin filaments are redistributed during cell infection with Dengue virus (DENV) as they are essential for the anchorage of the virus replication complex [25]. In uninfected HESC vimentin displayed a tipically extended network arrangement from the perinuclear region outwards to the cell plasma membrane (**Figure 2B**). Of note is the fact that ZIKV infection induced retraction of vimentin filaments from the cell periphery and their reorganization into dense structures in the perinuclear region (**Figure 2B**). Viral dsRNA IF showed a punctate structure that indeed co-localized with vimentin dense structures (**Figure 2B**).

Productive infection of two additional ZIKV strains, Puerto Rican 2015 (PRVABC59) and Thailand 2013 [26], was observed in primary HESC by both PFA and ddPCR of viral RNA (**Figure S1A** and**B,** respectively) 3 days post-infection, confirming intracellular co-localization of the E protein and dsRNA with calreticulin and vimentin, respectively (**Figure S1C** and **D**).

Thus, primary HESC are targets of *in vitro* infection and actively support the replication of different ZIKV strains in the ER and in association with vimentin filaments.

### In vitro decidualization of T-HESC cells upregulates ZIKV productive infection

In order to test whether decidualization could influence ZIKV infection, we infected the immortalized T-HESC cell line either in unstimulated conditions or following *in vitro* decidualization induced by progesterone and cAMP, as reported [27]. As shown by IF with both anti-dsRNA and anti-E protein mAbs, the MR766 strain productively infected T-HESC. A significant increase of the percentage of infected cells of *ca.* 2-fold was observed by IF staining of dT-HESC vs. control cells (**Figure 3A)**.

**Figure 3.**
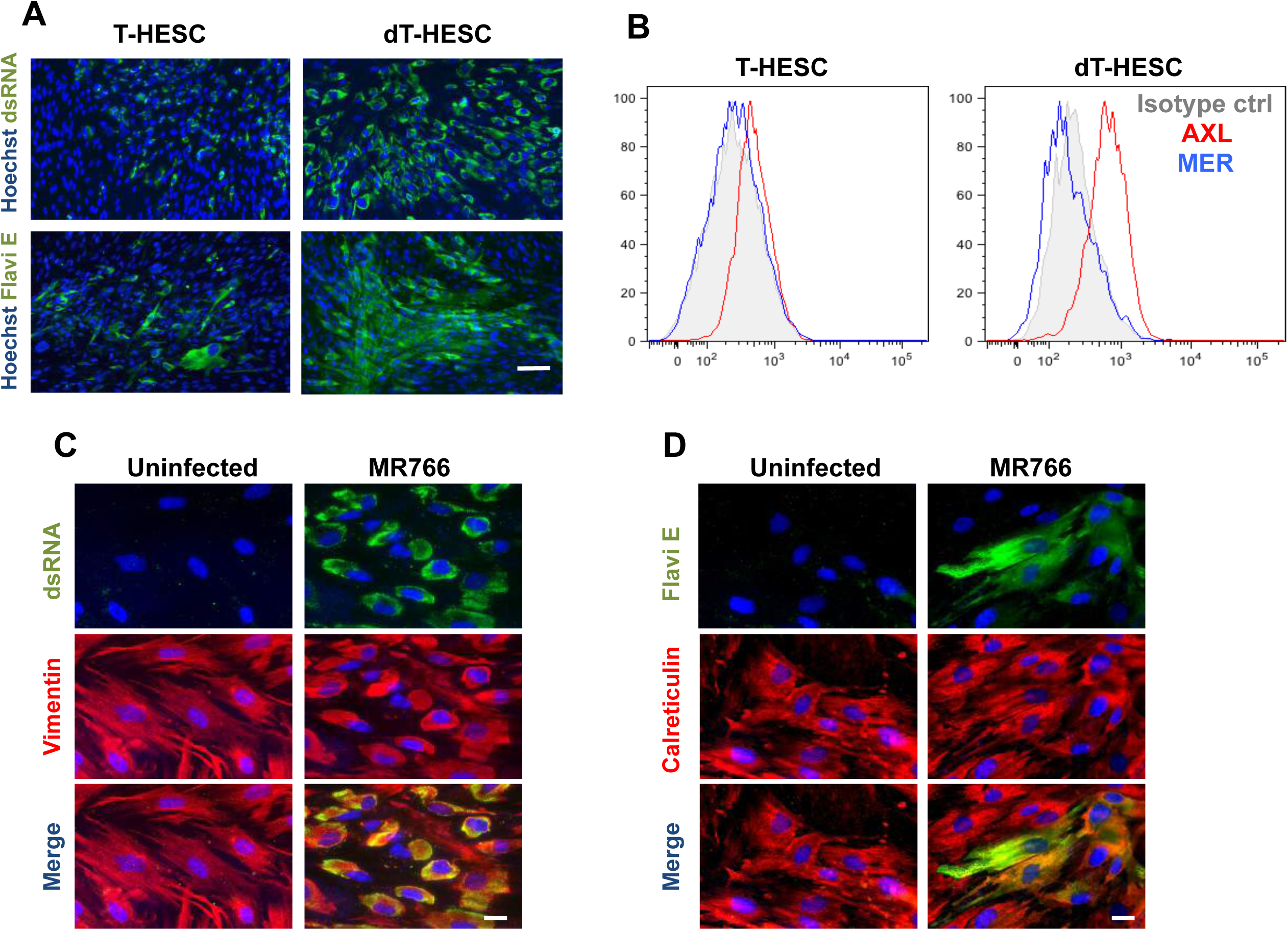
ZIKV infection of unstimulated and decidualized T-HESC cell line. **(A)** T-HESC (left) and decidualized (d)T-HESC (right) were stained with anti-dsRNA or Flavi E mAbs whereas the nuclei were stained with Hoechst. Scale bar: 20 µm. **(B)** Surface expression of AXL (red) and MER (blue) in T-HESC (left) and dT-HESC (right) was determined by flow cytometry. The histograms of one experiment representative of 3 independently performed are shown. Double immunostaining for Flavi E and calreticulin **(C)** or vimentin **(D)** in T-HESC (left) and dT-HESC (right) either uninfected or infected with MR766 at 72 h post-infection; Hoechst was used to stain nuclei. Scale bar: 10 µm.

The expression of two putative entry co-receptors for ZIKV entry, AXL and MER, was evaluated by cytofluorimetric analysis in both uninfected T-HESC and dT-HESC. While MER expression was not detectable, AXL was expressed by unstimulated T-HESC and, of note, it was upregulated in dT-HESC (RFI: 2.76 vs. 1.64, respectively; **Figure 3B**), consistently with the higher infection efficiency observed in these experimental conditions.

As observed with primary cells, calreticulin staining in uninfected cells appeared as a perinuclear fibrillar fluorescence whereas it expanded in the cytoplasm towards the cell perifery and colocalized with viral E protein expression in infected cells (**Figure 3C**). Punctate structures were observed in infected dT-HESCs stained with an anti-dsRNA Ab that colocalized with vimentin filaments in the perinuclear region (**Figure 3D**).

### ZIKV productive infection of T-HESC is cytopathic and induces the expression of IFN-β and ISGs

By PFA we next determined the kinetics of virus replication in T-HESC and dT-HESC cells by analyzing their culture supernatants in Vero cells. Virus replication in T-HESC of both MR766 and INMI-1 strains increased by approximately 10- and 5-fold over the levels of input virus, respectively, peaking around 96 h post-infection (**Figure 4A, left panel**). Decidualization of T-HESC increased the efficiency of virus replication by up to two orders of magnitude after 144 h post-infection (**Figure 4A, right panel**). Peak virus replication was detected by ddPCR 96 h post-infection and levels *ca.* 10 fold higher than those of the input virus were maintained up to 144 h post-infection (**Figure 4B**). ZIKV infection of both untreated T-HESC and dT-HESC induced cytopathicity and cell death as determined by the levels of adenylate kinase (AK) released in the culture supernatants [28] with kinetics overlapping those of virus replication (**Figure 4C**).

**Figure 4.**
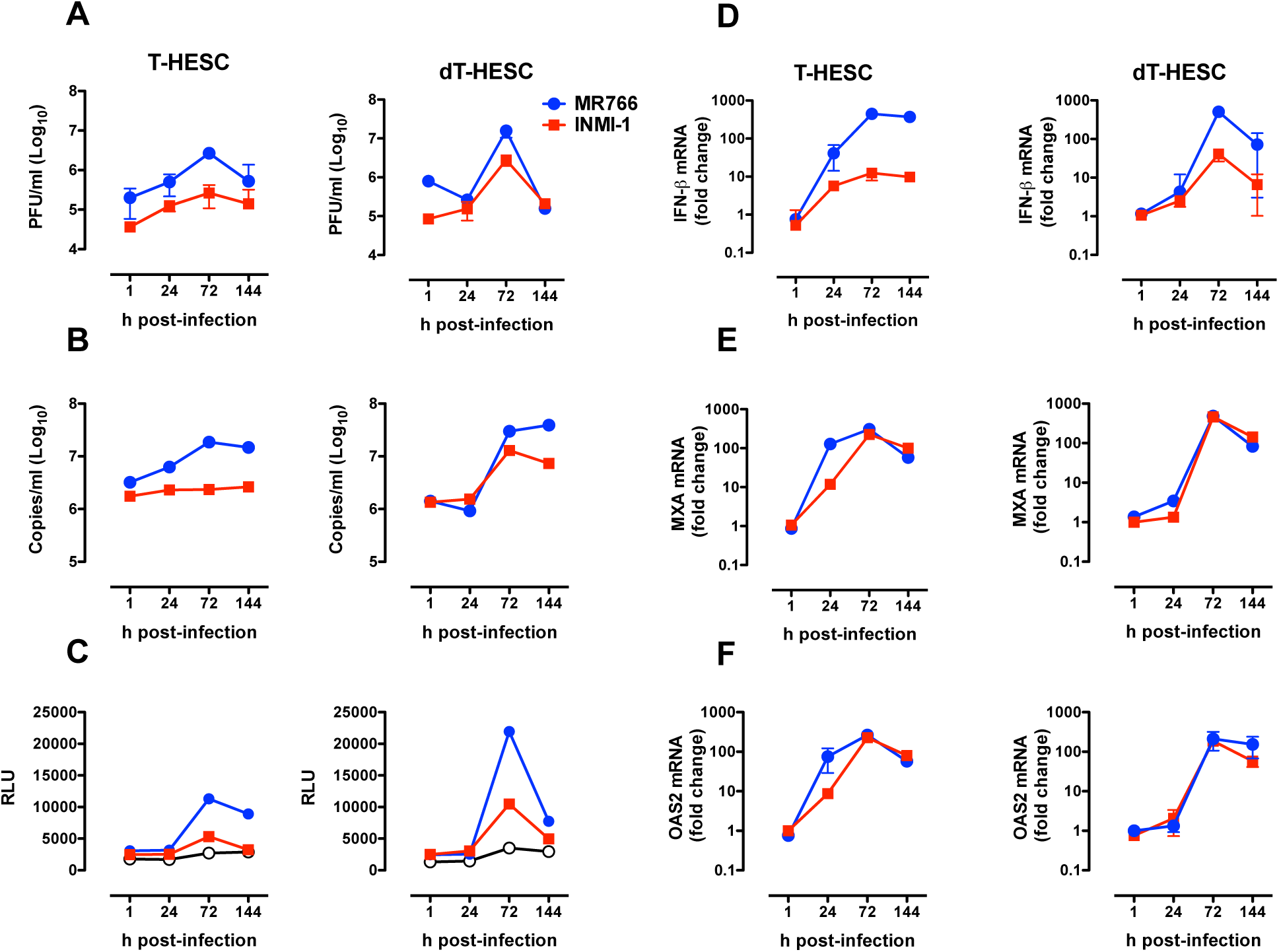
ZIKV replication, cytopathicity and induction of IFN-b and ISG expression in T-HESC and dT-HESC. Kinetics of virus replication in T-HESC (left) and dT-HESC (right) were measured by retrotitration of culture supernatants in Vero cells by PFA (**A**) and by ddPCR (**B**). (**C**) Kinetics of cell death were measured by the activity of cell-associated adenylate kinase released in cell culture supernatants. Open circles indicate uninfected cells. Time course of IFN-β mRNA (**D**), MXA (**E**) and OAS2 (**F**) expression in T-HESC (left) and dT-HESC (right) quantified by RT-qPCR. The results are expressed as the fold induction of transcripts in ZIKV-infected cells relative to those of uninfected cells. The results of one experiment representative of 3 independently conducted are shown.

Since it has been reported that ZIKV infection leads to the transcription and release of IFN-β in human skin fibroblasts [29], we evaluated the kinetic of IFN-β mRNA expression by RT-qPCR after infection of T-HESC. Indeed, higher levels of IFN-β gene transcription were detected 24 h after infection vs. those of uninfected cells and increased of ca. 300-fold 6 days after infection of T-HESC with the MR766 strain, whereas lower levels were induced upon infection with the INMI-1 strain (**Figure 4D**). The virus-induced levels of IFN-β mRNA in dT-HESC cells were similar to those of T-HESC cells; however, a more rapid insurgence of IFN-β expression was observed in T-HESC vs. dT-HESC (**Figure 4D**).

We next determined whether biologically active IFN-β was indeed secreted following ZIKV infection of T-HESC and dT-HESC cells. For this purpose, cell culture supernatants were tested on HEK-Blue IFN-α/β cells containing the secreted alkaline phosphatase (SEAP) reporter gene under the control of IFN-α/β inducible ISG54 promoter [30]. The induction of SEAP activity confirmed the presence of bioactive type 1 IFN in the supernatants of infected cells with moderately delayed kinetics in comparison to the upregulation of IFN-β mRNA observed in infected cells (**Figure S2**).

Among ISGs, myxovirus resistance protein 1 (MXA) and 2'-5'-Oligoadenylate Synthetase (OAS2) were previously shown to be genes that were mostly upregulated by ZIKV infection of fibroblasts [29]; therefore, we determined their kinetics of expression by RT-PCR. Both MXA and OAS2 mRNAs were upregulated in ZIKV infected vs. uninfected cells (**Figure 4E** and **F**, respectively) with kinetics almost superimposable to those of IFN-β mRNA expression; no quantitative differences were observed for these ISGs in cells infected with the MR766 and INMI-1 strains, in spite of different levels of virus replication.

## DISCUSSION

In the present study, we report for the first time that primary HESC are highly permissive to ZIKV productive infection by both an historic and different contemporary strains. We also showed that the T-HESC cell line, which models HESC in the proliferative phase, was permissive to ZIKV infection and supported virus replication. Infection of T-HESC strongly upregulated the expression of bioactive IFN-β and of related ISGs (MXA, OAS2). Interestingly, *in vitro* decidualization of T-HESC, increased the efficiency of ZIKV replication by up to 100-fold in association with the upregulation of the putative entry co-receptor AXL, but not of MER, suggesting a potential correlate of the increased levels of virus replication observed upon decidualization of the cell line.

Flaviviruses bind to a variety of surface molecules that serve as entry mediators or cofactors including the TAM family of tyrosine kinase receptors [31]. Among TAMs, AXL was reported to mediate ZIKV entry in dermal fibroblasts and epidermal keratinocytes [29]. Furthermore, AXL is highly expressed by neural stem cells, a privileged target of ZIKV infection in the fetal central nervous system [32] whereas it might play a minor role as compared with other TAM entry factors in placental cells [33]. Of importance is the observation that AXL was confirmed as ZIKV entry factor in a CRISPR/Cas9 screen of HeLa cells [34]. Our results support the hypothesis of a prominent role of AXL in infection of HESC and a potential deleterious role of progesterone in favoring virus transmission to the trophoblasts, in case of pregnancy, or from and to a male partner during sexual intercourse.

As for DENV, we observed that ZIKV dsRNA associated with vimentin intermediate filaments that were rearranged in the perinuclear region of infected HESC and T-HESC [25]. This observation suggests a potential role of vimentin in ZIKV RNA amplification machinery, perhaps by stabilization of replication complexes, as suggested for DENV replication [25]. We also observed that ZIKV E protein was localized in the ER of infected HESC and T-HESC cells, which showed dilated *cisternae* as a sign of massive expansion of the cell secretory machinery. These observations indicate that ZIKV, similarly to DENV, assembles and accumulates its progeny virions in the ER sacs [35] where a subset of ER-associated signal peptidase complex proteins is responsible for the proper cleavage of flavivirus structural proteins and secretion of viral particles [36]. The kinetics of ZIKV replication well correlated with the production of active IFN-β and the consequent induction of ISGs, such as MXA and OAS2, that likely curtailed spreading of infection.

Very recently, ZIKV has been detected in the FRT after the onset of clinical symptoms [37] and both male-to-female [15,38] and female-to-male transmission [20] have been documented suggesting that this vector-independent route of viral transmission might have occurred more frequently than reported. Furthermore, ZIKV presence in the FRT can foster vertical transmission from mother to fetus, as recently demonstrated in a mouse model of vaginal virus infection [13] and previously demonstrated for other members of the *Flaviviridae* family such as hepatitis C virus, in which vertical transmission from mother to child can occur in up to 10% of pregnancies [37]. As maternal decidual cells are in direct contact with extravillous trophoblast (EVT) on the tip of anchoring villi, one possibility is that ZIKV produced by infected HESC and/or other cells present in the FRT is transmitted to EVT in early pregnancy and then enters the fetal circulation.

An unclear aspect of ZIKV pathogenesis is whether ZIKV infection persists for longer in spite of its cytopathicity. If this is the case, ZIKV persistence in decidualized HESC might provide a viral reservoir for infection of cells of the *decidua basalis*, thus favoring viral transmission to chorionic villi.

Although ZIKV transmission occurs primarily through *Aedes* mosquitos bites, increasing evidence of vector-independent viral transmission through the sexual route bear potentially great societal implications for ZIKV spread [15]. Our *in vitro* findings indicate a vulnerability of the FRT to ZIKV infection, particularly upon endometrial decidualization by progesterone, as previously observed for other sexually transmitted viruses, such as herpes simplex virus and HIV [39]. As for these virus, topical microbicides developed and formulated for vaginal use [40] should be considered among the preventative strategies against ZIKV infection.

## METHODS

### Human Tissues

Endometrial biopsies from 7 pre-menopausal women with regular menstrual cycles (5 in proliferative phase and 2 in secretory phase) and from 1 perimenopause woman, all undergoing histeroscopy for diagnostic purposes, were obtained from the Ob&Gyn Department of the San Raffaele Scientific Institute in Milan. The selected women did not show any evident endometrial pathology or suffer from any endocrine disorder or systemic disease and have not received any steroid treatment for at least 3 months prior to tissue collection. Uterine samples were obtained by Vacuum Aspiration Biopsy Random Assay (Vabra). All women provided written informed consent before tissue collection and anonymized samples were delivered to the laboratory (see **Supporting Information** for the details of tissue culture preparation).

The immortalized human endometrial stromal cell line T-HESC, originated from normal human endometrial stromal cell immortalized with hTERT (Krikun et al, 2004), was obtained from ATCC (CRL-4003^TM^). Cells were maintained in complete growth medium according to the manufacturer’s instructions. To induce T-HESC cell decidualization, stromal cells were stimulated with cAMP (0.5 mM; Sigma) and medroxyprogesterone acetate (MPA) (1 μM; Sigma) for 7 days.

### ZIKV infection

The viral strain MR766 was obtained from the European virus archive (EVAg) and expanded in Vero cells. The Brazilian 2016/INMI-1 isolate (GenBank Accession # KU991811) was obtained from an Italian individual who traveled to Brazil in January 2016. HESC or T-HESC cells were seeded in 24 well plastic plates at 2.5x10^5^/ml; cell culture medium was removed from confluent cells and was replaced with virus containing supernatant at the MOI of 10. After 4 h, the supernatant was removed and fresh culture medium (0.5 ml) was added. The kinetics of virus replication were measured in supernatants collected 1, 24, 72 and 144 h post-infection and kept frozen at −80°C until further use (see **Supporting Information**).

### Indirect immunofluorescence of ZIKV E protein and dsRNA

Cells were fixed for 20 min in 4% paraformaldehyde solution (Sigma) in phosphate-buffered saline (PBS, Euroclone). Cells were permeabilized for 30 min in blocking solution, containing 0.2% Triton X-100 (Sigma) and 10% donkey serum (Sigma) and incubated overnight at 4°C with the primary mAb in blocking solution. The following mAbs specific for Flavivirus E protein (1:200, Millipore, MAB10216), double-stranded RNA (1:300, English and Scientific Consulting Kft, Hungary), vimentin (1:300, Bioss, BS-0756R) and calreticulin (1:300, Sigma, C4606) were used. Cells were then washed with PBS and incubated for 1h with Hoechst and either anti-mouse Alexa Fluor-488 or anti-rabbit Alexa Fluor-594 secondary Abs (1:1,000 in blocking solution, ThermoFisher Scientific). After PBS washes, cells were washed again and mounted.

### Flow Cytometry

Mouse anti-human AXL mAb (clone # 108724, MAB154) and mouse anti-human MER mAb (clone # 125518, MAB8912) was purchased from R&D Systems. Goat anti-mouse IgG secondary antibody, RPE conjugated was purchased from ThermoFisher Scientific (P-852). 2x10^4^ T-HESC were used per condition. Samples were acquired on FACSCanto (BD) flow cytometer. Dead cells were excluded on the basis of propidium iodide and DAPI staining. All data were analyzed using FlowJo software (Tree Star). Relative fluorescent intensity (RFI) of AXL and MER expression on T-HESC was calculated dividing the mean AXL and MER fluorescence intensity by the mean fluorescence intensity of the corresponding isotype control mAb.

### Cell death detection assay

10 μl samples of culture supernatant were transferred on a half black 96 well plate (Costar). To each well, 50 μl of the adenylate kinase detection reagent (ToxiLight^®^ BioAssay, Lonza) was added and the plate was incubated for 10 min at room temperature. Luminescence was measured in a Mithras LB940 Microplate Reader (Berthold Technologies). The results were expressed as relative luminescent unit (RLU).

### RT-qPCR for IFN-β and ISGs

Total RNA was extracted from either non-decidualized or decidualized T-HESC cells by using a TRIzol Plus RNA purification kit, followed by DNase I treatment (Invitrogen). cDNA was synthesized from total RNA (1 μg) using a SuperScript first-strand synthesis system (Invitrogen) with random hexamers. SYBR green (Applied Biosystems) qPCR was performed with 50 ng of cDNA in a total volume of 25 µl with the primer pair (250 nM) described in the **Table S1**. All reactions were performed with an ABI 7700 Prism instrument (Applied Biosystems). mRNA expression was calculated by using the relative quantification method to uninfected samples after normalized to human glyceraldehyde-3-phosphate dehydrogenase (GAPDH) mRNA expression.

### Statistical Analysis

Prism GraphPad software v. 4.0 (www.graphpad.com) was used for all statistical analyses. Comparison between two groups was performed using the Student’s t-test.

## ACKNOWLEDGMENTS

We thank Maria Rosaria Capobianchi, National Institute for Infectious Diseases “Lazzaro Spallanzani”, Rome, Italy and Edwin Yates, University of Liverpool, UK for critical reading of the manuscript and helpful suggestions.

## SUPPORTING INFORMATION CAPTIONS

**Fig S1**. Infection of HESC with Puerto Rico 2015 and Thailand 2013 ZIKV strains.

**Fig S2**. Time course of bioactive IFN-β release in response to ZIKV in T-HESC and dT-HESC.

**Table S1**. Primers used in the study

Supporting experimental procedures

## REFERENCES

1. Musso D, Gubler DJ (2016) Zika Virus. Clinical microbiology reviews 29: 487–524.

2. Dick GW, Kitchen SF, Haddow AJ (1952) Zika virus. I. Isolations and serological specificity. Transactions of the Royal Society of Tropical Medicine and Hygiene 46: 509–520.

3. Cao-Lormeau VM, Blake A, Mons S, Lastere S, Roche C, et al. (2016) Guillain-Barre Syndrome outbreak associated with Zika virus infection in French Polynesia: a case-control study. Lancet 387: 1531–1539.

4. Besnard M, Eyrolle-Guignot D, Guillemette-Artur P, Lastere S, Bost-Bezeaud F, et al. (2016) Congenital cerebral malformations and dysfunction in fetuses and newborns following the 2013 to 2014 Zika virus epidemic in French Polynesia. Euro surveillance: bulletin Europeen sur les maladies transmissibles = European communicable disease bulletin 21.

5. Brasil P, Pereira JP, Jr., Raja Gabaglia C, Damasceno L, Wakimoto M, et al. (2016) Zika Virus Infection in Pregnant Women in Rio de Janeiro - Preliminary Report. The New England journal of medicine.

6. Driggers RW, Ho CY, Korhonen EM, Kuivanen S, Jaaskelainen AJ, et al. (2016) Zika Virus Infection with Prolonged Maternal Viremia and Fetal Brain Abnormalities. The New England journal of medicine.

7. Calvet G, Aguiar RS, Melo AS, Sampaio SA, de Filippis I, et al. (2016) Detection and sequencing of Zika virus from amniotic fluid of fetuses with microcephaly in Brazil: a case study. The Lancet Infectious diseases.

8. Adibi JJ, Marques ET, Jr., Cartus A, Beigi RH (2016) Teratogenic effects of the Zika virus and the role of the placenta. Lancet 387: 1587–1590.

9. Bayer A, Lennemann NJ, Ouyang Y, Bramley JC, Morosky S, et al. (2016) Type III Interferons Produced by Human Placental Trophoblasts Confer Protection against Zika Virus Infection. Cell host & microbe 19: 705–712.

10. Quicke KM, Bowen JR, Johnson EL, McDonald CE, Ma H, et al. (2016) Zika Virus Infects Human Placental Macrophages. Cell host & microbe 20: 83–90.

11. Tabata T, Petitt M, Puerta-Guardo H, Michlmayr D, Wang C, et al. (2016) Zika Virus Targets Different Primary Human Placental Cells, Suggesting Two Routes for Vertical Transmission. Cell host & microbe.

12. Miner JJ, Cao B, Govero J, Smith AM, Fernandez E, et al. (2016) Zika Virus Infection during Pregnancy in Mice Causes Placental Damage and Fetal Demise. Cell 165: 1081–1091.

13. Yockey LJ, Varela L, Rakib T, Khoury-Hanold W, Fink SL, et al. (2016) Vaginal Exposure to Zika Virus during Pregnancy Leads to Fetal Brain Infection. Cell.

14. D’Ortenzio E, Matheron S, Yazdanpanah Y, de Lamballerie X, Hubert B, et al. (2016) Evidence of Sexual Transmission of Zika Virus. The New England journal of medicine 374: 2195–2198.

15. Grischott F, Puhan M, Hatz C, Schlagenhauf P (2016) Non-vector-borne transmission of Zika virus: A systematic review. Travel Medicine and Infectious Disease.

16. Nicastri E, Castilletti C, Liuzzi G, Iannetta M, Capobianchi MR, et al. (2016) Persistent detection of Zika virus RNA in semen for six months after symptom onset in a traveller returning from Haiti to Italy, February 2016. Euro surveillance: bulletin Europeen sur les maladies transmissibles = European communicable disease bulletin 21.

17. Atkinson B, Hearn P, Afrough B, Lumley S, Carter D, et al. (2016) Detection of Zika Virus in Semen. Emerging infectious diseases 22: 940.

18. Musso D, Roche C, Robin E, Nhan T, Teissier A, et al. (2015) Potential sexual transmission of Zika virus. Emerging infectious diseases 21: 359–361.

19. Mansuy JM, Pasquier C, Daudin M, Chapuy-Regaud S, Moinard N, et al. (2016) Zika virus in semen of a patient returning from a non-epidemic area. The Lancet Infectious diseases 16: 894–895.

20. Davidson A, Slavinski S, Komoto K, Rakeman J, Weiss D (2016) Suspected Female-to-Male Sexual Transmission of Zika Virus - New York City, 2016. MMWR Morbidity and mortality weekly report 65: 716–717.

21. Brosens JJ, Gellersen B (2006) Death or survival--progesterone-dependent cell fate decisions in the human endometrial stroma. Journal of molecular endocrinology 36: 389–398.

22. Blazquez AB, Escribano-Romero E, Merino-Ramos T, Saiz JC, Martin-Acebes MA (2014) Stress responses in flavivirus-infected cells: activation of unfolded protein response and autophagy. Frontiers in microbiology 5: 266.

23. Aebi M, Bernasconi R, Clerc S, Molinari M (2010) N-glycan structures: recognition and processing in the ER. Trends in biochemical sciences 35: 74–82.

24. Osteen KG, Hill GA, Hargrove JT, Gorstein F (1989) Development of a method to isolate and culture highly purified populations of stromal and epithelial cells from human endometrial biopsy specimens. Fertility and sterility 52: 965–972.

25. Teo CS, Chu JJ (2014) Cellular vimentin regulates construction of dengue virus replication complexes through interaction with NS4A protein. Journal of virology 88: 1897–1913.

26. Fonseca K, Meatherall B, Zarra D, Drebot M, MacDonald J, et al. (2014) First case of Zika virus infection in a returning Canadian traveler. The American journal of tropical medicine and hygiene 91: 1035–1038.

27. Krikun G, Mor G, Alvero A, Guller S, Schatz F, et al. (2004) A novel immortalized human endometrial stromal cell line with normal progestational response. Endocrinology 145: 2291–2296.

28. Single B, Leist M, Nicotera P (1998) Simultaneous release of adenylate kinase and cytochrome c in cell death. Cell death and differentiation 5: 1001–1003.

29. Hamel R, Dejarnac O, Wichit S, Ekchariyawat P, Neyret A, et al. (2015) Biology of Zika Virus Infection in Human Skin Cells. Journal of virology 89: 8880–8896.

30. Frumence E, Roche M, Krejbich-Trotot P, El-Kalamouni C, Nativel B, et al. (2016) The South Pacific epidemic strain of Zika virus replicates efficiently in human epithelial A549 cells leading to IFN-beta production and apoptosis induction. Virology 493: 217–226.

31. Perera-Lecoin M, Meertens L, Carnec X, Amara A (2014) Flavivirus entry receptors: an update. Viruses 6: 69–88.

32. Nowakowski TJ, Pollen AA, Di Lullo E, Sandoval-Espinosa C, Bershteyn M, et al. (2016) Expression Analysis Highlights AXL as a Candidate Zika Virus Entry Receptor in Neural Stem Cells. Cell stem cell 18: 591–596.

33. Tabata T, Petitt M, Puerta-Guardo H, Michlmayr D, Wang C, et al. (2016) Zika Virus Targets Different Primary Human Placental Cells, Suggesting Two Routes for Vertical Transmission. Cell host & microbe 20: 155–166.

34. Savidis G, McDougall William M, Meraner P, Perreira Jill M, Portmann Jocelyn M, et al. (2016) Identification of Zika Virus and Dengue Virus Dependency Factors using Functional Genomics. Cell reports 16: 232–246.

35. Welsch S, Miller S, Romero-Brey I, Merz A, Bleck CK, et al. (2009) Composition and three-dimensional architecture of the dengue virus replication and assembly sites. Cell host & microbe 5: 365–375.

36. Zhang R, Miner JJ, Gorman MJ, Rausch K, Ramage H, et al. (2016) A CRISPR screen defines a signal peptide processing pathway required by flaviviruses. Nature 535: 164–168.

37. Prisant N, Bujan L, Benichou H, Hayot PH, Pavili L, et al. (2016) Zika virus in the female genital tract. The Lancet Infectious diseases.

38. Foy BD, Kobylinski KC, Chilson Foy JL, Blitvich BJ, Travassos da Rosa A, et al. (2011) Probable non-vector-borne transmission of Zika virus, Colorado, USA. Emerging infectious diseases 17: 880–882.

39. Wira CR, Rodriguez-Garcia M, Patel MV (2015) The role of sex hormones in immune protection of the female reproductive tract. Nature reviews Immunology 15: 217–230.

40. Alexandre KB, Mufhandu HT, London GM, Chakauya E, Khati M (2016) Progress and Perspectives on HIV-1 microbicide development. Virology 497: 69–80.

